# Flowering behaviour in *Arabis alpina* ensures the maintenance of a perennating dormant bud bank

**DOI:** 10.1101/562868

**Authors:** Alice Vayssières, Priyanka Mishra, Adrian Roggen, Udhaya Ponraj, Ulla Neumann, Klaus Theres, Karin Ljung, Maria C. Albani

## Abstract

*Arabis alpina*, similar to woody perennials, has a complex architecture with a zone of axillary vegetative branches and a zone of dormant buds that serve as perennating organs. We show that floral development during vernalization is the key for shaping the dormant bud zone by facilitating a synchronized and rapid growth after vernalization and thereby causing an increase in auxin response and transport and endogenous indole-3-acetic acid (IAA) levels in the stem. Floral development during vernalization is associated with the development of axillary buds in subapical nodes. Our transcriptome analysis indicated that these buds are not dormant during vernalization but only attain sustained growth after the return to warm temperatures. Floral and subapical vegetative branches grow after vernalization and inhibit the development of the buds below. Dormancy in these buds is regulated across the *A. alpina* life cycle by low temperatures and by apical dominance in a BRANCHED 1-dependent mechanism.

## INTRODUCTION

Bud dormancy plays an important role in survival through harsh environmental conditions and long-term growth^1^. Thus, during the perennial life cycle, axillary and apical buds transition through the various stages of dormancy before they resume active organogenesis and develop into flowering or vegetative branches^2^. During development the outgrowth of axillary buds close to the shoot apical meristem is repressed by apical dominance, a phenomenon also known as correlative inhibition, latency or paradormancy, which occurs in both annual and perennial species^3,4^. This form of dormancy is not definitive and paradormant buds can resume growth when the main shoot apex is removed^5^. Buds in trees and herbaceous perennials also enter two other forms of dormancy, endo- and eco-dormancy^2^. Endodormancy is regulated by endogenous signals within the bud whereas ecodormancy is imposed by unfavorable environmental conditions^6^. During the life cycle of a perennial plant, apical or axillary buds experience winter in the endodormant state and later become ecodormant so that they will actively grow only during favorable environmental conditions. It is however very common in perennials for branches to have axillary buds during spring and summer, which later on will stay dormant across multiple growing seasons. These dormant buds serve as a backup bud bank and, in case of damage, are used as reservoirs for potential growth facilitating a bet-hedging mechanism^7,8^. Interestingly, dormant buds and actively growing (vegetative or reproductive) axillary branches are organized in zones in a species specific pattern^9,10^.

The outgrowth of an axillary bud after decapitation involves two phases, firstly the rapid release from dormancy and secondly its sustained growth^11^. Auxin, strigolactones, cytokinin and sugar fine-tune this process by regulating the expression of the TCP transcription factor BRANCHED 1 (BRC1)^12^. Decapitation causes an elevation of sucrose levels followed by a depletion of the endogenous indole-3-acetic acid (IAA) in the polar auxin transport stream, which influences the auxin flux out of the axillary bud contributing to its sustainable growth^13–17^. Flowering transition triggers the activation of the upper most axillary buds in a basipetal sequence having a similar effect to decapitation^13,18^. In an annual plant, the relation between flowering and bud activation is easy to trace as its life cycle ends within one growing season. However, many perennials spread the flowering process over several years and floral bud initiation is temporally separated from anthesis.

Here we used the perennial *Arabis alpina,* a close relative of *Arabidopsis thaliana*, to investigate the link between flowering and the maintenance of a dormant bud bank in perennials. The shoot architecture of *A. alpina* is polycarpic and consists of branches that undergo flowering, others that remain vegetative and nodes with dormant axillary buds^19^. *A. alpina* mutants or transgenic lines with reduced function in components of the age pathway influence the fate of vegetative branches but still require exposure to prolonged cold to flower, a process known as vernalization^20–23^. Interestingly, the maintenance of dormant buds is only compromised in mutants that do not require vernalization^19^. The most described example is the *perpetual flowering 1* (*pep1*) mutant which carries lesions in the ortholog of the MADS box transcription factor FLOWERING LOCUS C (FLC)^19,24,25^. Here we show that the requirement of vernalization to flower is linked to the maintainance of dormant axillary buds.

## RESULTS

### Flowering during vernalization correlates with the formation of a perennial shoot architecture including a zone of dormant axillary buds

To assess the relationship between flowering and plant architecture, we exposed the accession Pajares to different durations of vernalization that influence flowering and we scored bud activity and fate. As shown previously, in 8-week-old plants vernalized for 12 weeks the shoot apical meristem (I zone, nodes 29–40) and the lower axillary branches (V1 zone, nodes 1–12) flowered (Fig. 1a,e,f)^10,19^. The architecture of *A. alpina* plants was already established 3 weeks after vernalization, showing a clear inhibition of axillary bud growth in nodes 12–21, with leaf axils showing no visible outgrowth or buds that did not grow more than 0.5 cm (V2 zone, Fig. 1a,b,e,f)^10^. The nodes above (nodes 22–29) contained buds that actively grew after vernalization and giving rise to axillary branches that remained vegetative (V3 zone, Fig. 1b,e, f)^10,19,26^. When we compared the architecture of plants after vernalization with the one before vernalization we observed a similar pattern of growth as described previously^19,26,27^. The V1 branches developed before vernalization, the V2 buds were present on the axils of leaves before vernalization, and the V3 branches arose in the axils of leaves that grew during vernalization. It has been recently shown that extended vernalization is required to ensure floral commitment^10^. To test whether extended vernalization influences the size of the V2 zone, we exposed 8-week-old plants to 12, 15, 18, 21 and 24 weeks of vernalization (Supplementary Fig. 1). The outgrowth of V3 branches and the inflorescence stem was accelerated after longer durations of vernalization (Supplementary Fig. 1a,b)^10^. However, we observed no difference in the number of dormant buds suggesting that extended vernalization does not have an impact on the V2 buds (Supplementary Fig. 1c,d). We then exposed plants to durations of vernalization shorter than 12 weeks. As shown previously, plants grown continuously in LDs or vernalized only for 3 weeks did not flower, whereas plants vernalized for 8 weeks showed extreme floral reversion phenotypes (Supplementary Fig. 2)^10,19^. To follow bud activity, we measured the length of axillary branches at each leaf axil 3 weeks after vernalization or after 11 weeks in LDs for non-vernalized plants. Branch length in LD grown plants and in 3 week vernalized plants was reduced acropetally (Fig. 1g,h). Interestingly, nodes 12–18 were completely inhibited only in plants exposed to 12 weeks of vernalization (Fig. 1g,h). In plants vernalized for 8 weeks, the zonation was visible by measuring the branch length (Fig. 1h) but no dormant bud zone was observed (Fig. 1g,h). Nevertheless, in both 8 week and 12 week vernalized plants the nodes occupied with actively growing branches were always just above the nodes with the inhibited buds suggesting that the presence of V3 branches might be correlated with the stable inhibition of bud growth in the V2 zone.

**Fig. 1:**
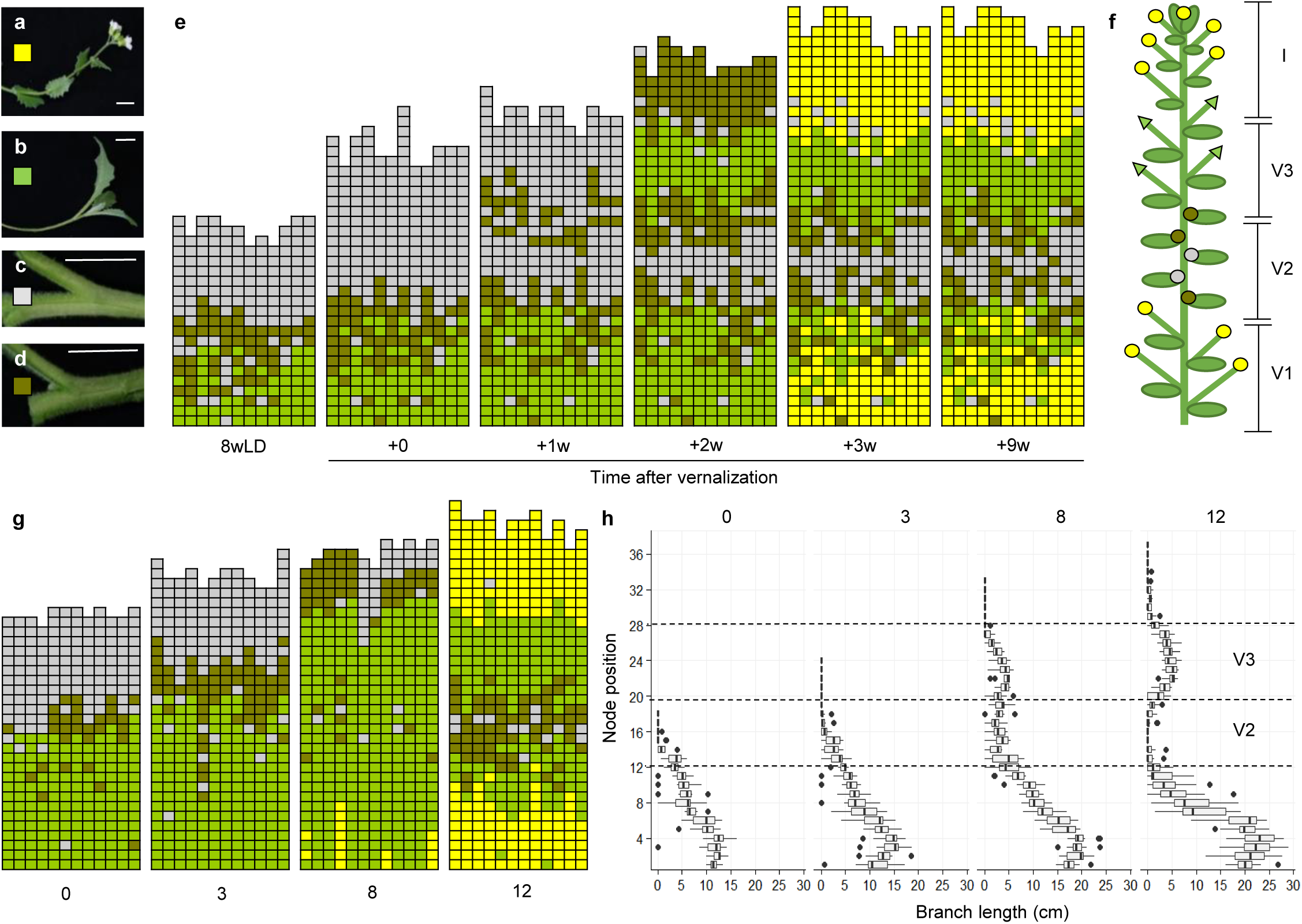
Flowering in the main shoot correlates with perennial shoot architecture in *A. alpina.* **a-e,** Analysis of branch formation in plants vernalized for 12 weeks which is sufficient for flowering in *A. alpina*. Plants were grown in long days (LDs) for 8 weeks (8wLD), vernalized for 12 weeks (+0) and transferred back to LDs for 1, 2, 3 and 9 weeks (+1w, +2w, +3w, +9w). Flowering plants have a complex plant architecture and some axillary branches flower (**a**) or grow vegetatively (**b**). Flowering plants also have leaf axils that are empty (**c**) or contain a branch smaller than 0.5 cm (**d**). In (**e**) each column represents a single plant and each square within a column indicates an individual leaf axil. The bottom row represents the oldest leaf axil. *Yellow* denotes the presence of a flowering branch (**a**). *Green* denotes the presence of a vegetative axillary branch (**b**). *Grey* denotes an empty leaf axil (**c**). *Brown* denotes a leaf axil with a branch smaller than 0.5 cm (**d**). **f**, This plant architecture is organized in zones described as V1, V2, V3 and I in Lazaro et al.^10^. *Yellow* circle denotes flowering of the main or side shoots, *grey* and *brown* circle indicate the presence of dormant buds (V2), *green* triangle the presence of a vegetative branch. **g**, Analysis of branch formation and **h**, branch length in *A. alpina* plants exposed to shorter lengths of vernalization that do not secure flowering. Plants were grown for 11 weeks in LDs (0), or for 8 weeks in LDs and subsequently vernalized for 3 (3w), 8 (8w) or 12 (12w) weeks. Plants were scored 3 weeks after they were returned to a LD greenhouse, *n*=12. Bar size indicates 1 cm.

*PEP1* determines the requirement for vernalization to flower and the fate of the V3 branches after vernalization^10,19^. To test whether *PEP1* plays a role in the zonation of *A. alpina* shoots, we measured branch length and the fate of the axillary branches in the *pep1-1* mutant compared to the wild-type (Supplementary Fig. 3). *pep1-1* flowered after 8 weeks in LDs and showed a deviation in branching phenotype compared to the wild-type (Supplementary Fig. 3). Nevertheless, both genotypes lacked the characteristic zonation observed in flowering plants after vernalization suggesting that flowering in LDs does not ensure perennial plant architecture in *A. alpina*. To test whether flowering during vernalization correlates with the repression of V2 buds, we exposed plants of different ages to vernalization. Earlier reports showed that 3-week-old plants are not competent to flower in response to vernalization and remain vegetative, whereas 5-week-old plants flower (Supplementary Fig. 4)^20,21,26^. Plants grown for 3 weeks prior to vernalization did not show bud inhibition at any node below the newly formed branches, whereas in 5-week-old plants they did in nodes 3–8 (Supplementary Fig. 4c). These plants also contained V3 vegetative branches above the inhibited buds (nodes 8–19) (Supplementary Fig. 4c). Altogether, these results suggest that zones of differential bud activity in *A. alpina* are formed only in genotypes that initiate flowering during vernalization.

### V3 buds are not dormant at the end of vernalization, but attain sustained growth only after the return to warm temperatures

In *A. thaliana* and other species, bud growth has been demonstrated using the auxin inducible synthetic promoter *DR5*^13,17^. To check whether V2 and V3 buds are active during or after vernalization, we developed transgenic *A. alpina* plants carrying the *DR5* promoter fused to the reporter gene β*-glucuronidase* (*GUS*). We observed the GUS signal only after vernalization in the vasculature of V3 buds and in the main stem just below the leaf nodes (Fig. 2 a-d). These data suggest that at 5 days after vernalization the V3 buds are growing whereas the V2 buds are not.

**Fig. 2:**
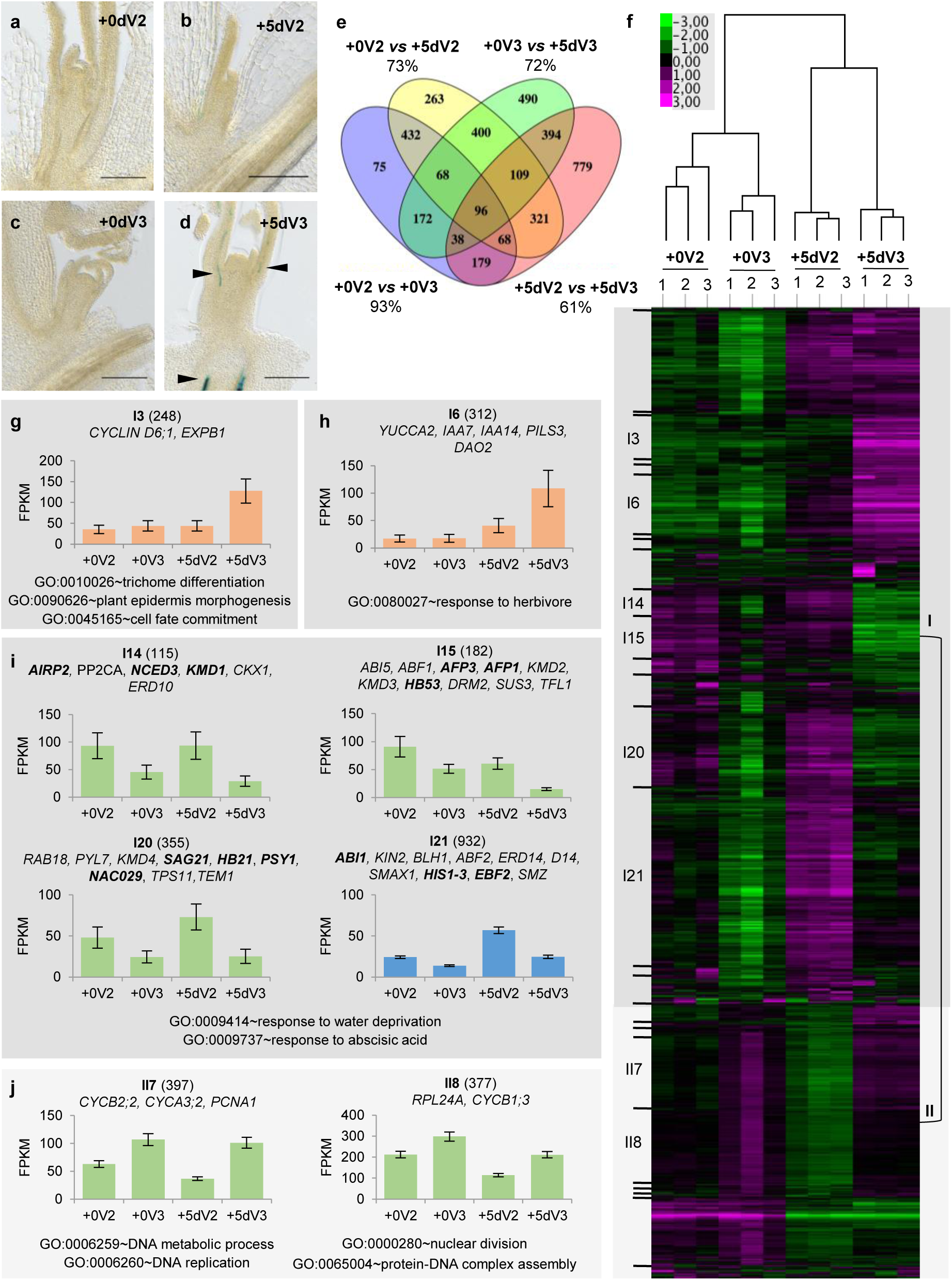
The transcriptome of buds that will develop into axillary vegetative branches or stay dormant is already differentiated during vernalization. **a-d**, GUS activity in longitudinal sections of axillary buds harvested from the nodes that will stay dormant (V2; **a** and **b**) or give rise to axillary vegetative branches (V3; **c** and **d**) at the end of vernalization (+0; **a** and **c**) and five days after vernalization (+5d; **b** and **d**) in *DR5::GUS A. alpina* plants. *Arrows* indicate GUS signal. **e**, Venn diagram represents the overlap of genes significantly regulated in the four comparisons tested. Percentages indicate overlap with differentially expressed genes between comparisons. **f**, Heat map representing the hierarchical clustering of 4983 coexpressed transcripts between V2 and V3 buds at the end of vernalization (+0) and 5 days after vernalization (+5d). Coexpressed clusters were assigned into two higher level clusters (I and II). **g-j** Selected clusters with common GOs are shown as FPKM (+/− SE) values. Colors indicate specific pattern of interest; *orange*: genes upregulated specifically in V3 buds after vernalization, *blue*: genes upregulated specifically in V2 buds after vernalization, *green*: genes differentially expressed between V2 and V3 buds at the end of vernalization. Numbers in brackets indicate the number of genes present in each cluster. Gene names indicated in bold have been annotated as “ *bud dormancy*” genes in Tarancon et al.^39^.

To investigate the molecular mechanisms that lead to the activation of V3 buds and the inhibition of growth in V2 buds, we performed a transcriptome profiling on dissected buds directly at the end of vernalization and 5 days after vernalization. The transcriptome of V2 and V3 buds at 5 days after vernalization was the most dissimilar (1984 genes; +5dV2 *vs* +5dV3; Fig. 2e; Supplementary Table 1). However, the transcriptome of V2 and V3 buds differed also at the end of vernalization (1128 genes for +0V2 *vs* +0V3; Fig. 2e; Supplementary Table 1). This suggests that the V2 and V3 buds were differentiated already during vernalization although active growth of the V3 buds was only observed after vernalization. Interestingly, 93% of the genes differentially expressed between V2 and V3 buds at the end of vernalization were differentially expressed between samples after vernalization or/and between developmental stages (Fig. 2e). Likewise, Gene Ontology (GO) enrichment analysis indicated that all GOs enriched for the differentially expressed genes between V2 and V3 buds at the end of vernalization were also found in the other comparisons (Supplementary Fig. 5, Supplementary Table 2). GO terms common to all comparisons were mainly associated with hormone responses such as abscisic acid, ethylene and jasmonic acid (Supplementary Fig. 5). These results suggest that hormones play an important role in the activation of V3 buds and the repression of V2 buds. To identify genes that share similar expression patterns, we performed hierarchical clustering analysis. We identified 34 co-expressed clusters, which were assigned into two higher level clusters I and II (Fig. 2f). Interestingly, the separation of these higher level clusters was shaped by the expression of genes in the V3 buds during vernalization. Genes in Cluster I showed low and genes in Cluster II showed high transcript accumulation in V3 buds at the end of vernalization (Fig. 2f). Genes in Cluster I3 and I6 showed higher transcript accumulation in V3 buds after vernalization accounting for putative candidate genes involved in the sustained growth of V3 buds (Fig. 2g,h). Specifically, in Cluster I3 most enriched GO terms among the identified genes were associated with developmental processes including genes involved in cell expansion (*EXPB1*) (Fig. 2g, I3, Supplementary Table 3). This result clearly supports that V3 buds only grow after vernalization. The growth V3 buds was also associated with the upregulation of transcript accumulation of the *YUCCA2* homolog, a gene coding for an enzyme that catabolizes the biosynthesis of IAA in *A. thaliana*, and the auxin signaling factors *IAA7* and *IAA14*, whose expression levels reflect also auxin levels, identified in cluster I6 (Fig. 2h)^28,29^. These results confirm our *DR5* results that the activation of auxin response occurs in V3 buds after vernalization. Clusters I14, I15, I20, I21, II7 and II8 showed different transcript accumulation between buds already during vernalization (Fig. 2i,j). Genes in these clusters also showed similar differences in transcript accumulation after vernalization indicating that the majority of the differences observed during vernalization are also maintained afterwards. Genes in Clusters II7 and II8 showed higher transcript accumulation in V3 buds compared to V2 buds and were enriched for GO terms related to cell division, which included homologs of cell cycle regulators, such as *PCNA1,* previously shown to be upregulated during bud activation in other species (Fig. 2j, Supplementary Fig. 3)^30,31^. These results suggest that genes involved in cell cycle and transcription machinery are upregulated in V3 buds during vernalization. Clusters I14, I15, I20 and I21 showed lower transcript accumulation in V3 buds compared to V2 buds. Interestingly, all these clusters were enriched for GOs related to response to abscisic acid and water deprivation and contained genes related to ABA signaling (e.g. *AIRP2, ABF1*, *ABF2*, *ABI1, ABI5, KIN2*, *AFP1 AFP3, BLH1* and *PP2CA*) or ABA biosynthesis (e.g. *NCED3*) shown to be associated with bud dormancy in several species (Fig. 3i)^32–35^. We also detected the dehydrin coding genes *ERD10*, *ERD14* and *RAB18*, which are induced by ABA and suggested to prevent water dehydration during tree winter dormancy (Fig. 3i)^36^. Transcript accumulation of homologs of genes associated with repression of the cytokinin level and response such as *CKX1* and *KMD1–4* were also detected in these clusters (Fig. 3i)^37,38^. These data suggest that V3 buds during vernalization might contain lower levels of ABA that represses bud outgrowth and higher levels of cytokinin that promotes bud outgrowth compared to V2 buds. We also identified the homologs of genes that have been previously shown in *A. thaliana* to respond to conditions that trigger dormancy and therefore are considered as dormancy markers^39^. This includes the *A. alpina* homologs of *HB21*, *HB53*, *PSY1*, *NAC029*, *SAG21* and *HIS1-3* (Fig. 2i, Supplementary Table 5). Interestingly, genes in I21, in contrast to clusters I14, I15 and I20, showed a high transcript upregulation in V2 buds after vernalization (Fig. 2i). In addition to ABA signalling genes described before, we also detected the homologs of the strigolactone signaling genes *D14* and *SMAX1* in this cluster. D14 in *A. thaliana* and its homologs in other species regulate bud dormancy^40,41^. These results suggest that the dormancy status of the V2 buds is enhanced after vernalization which correlates with the activation of genes involved in ABA and SL signaling. Interestingly, we also identified homologs of the floral repressors such as *TEM1*, *TFL1* and *SMZ* that regulate flowering through the age pathway (Fig. 2i)^20,42–45^. This result suggests that V3 buds are more competent to flower compared to V2 buds during and after vernalization and may relate to the low dormancy status of these axillary meristems.

**Fig. 3:**
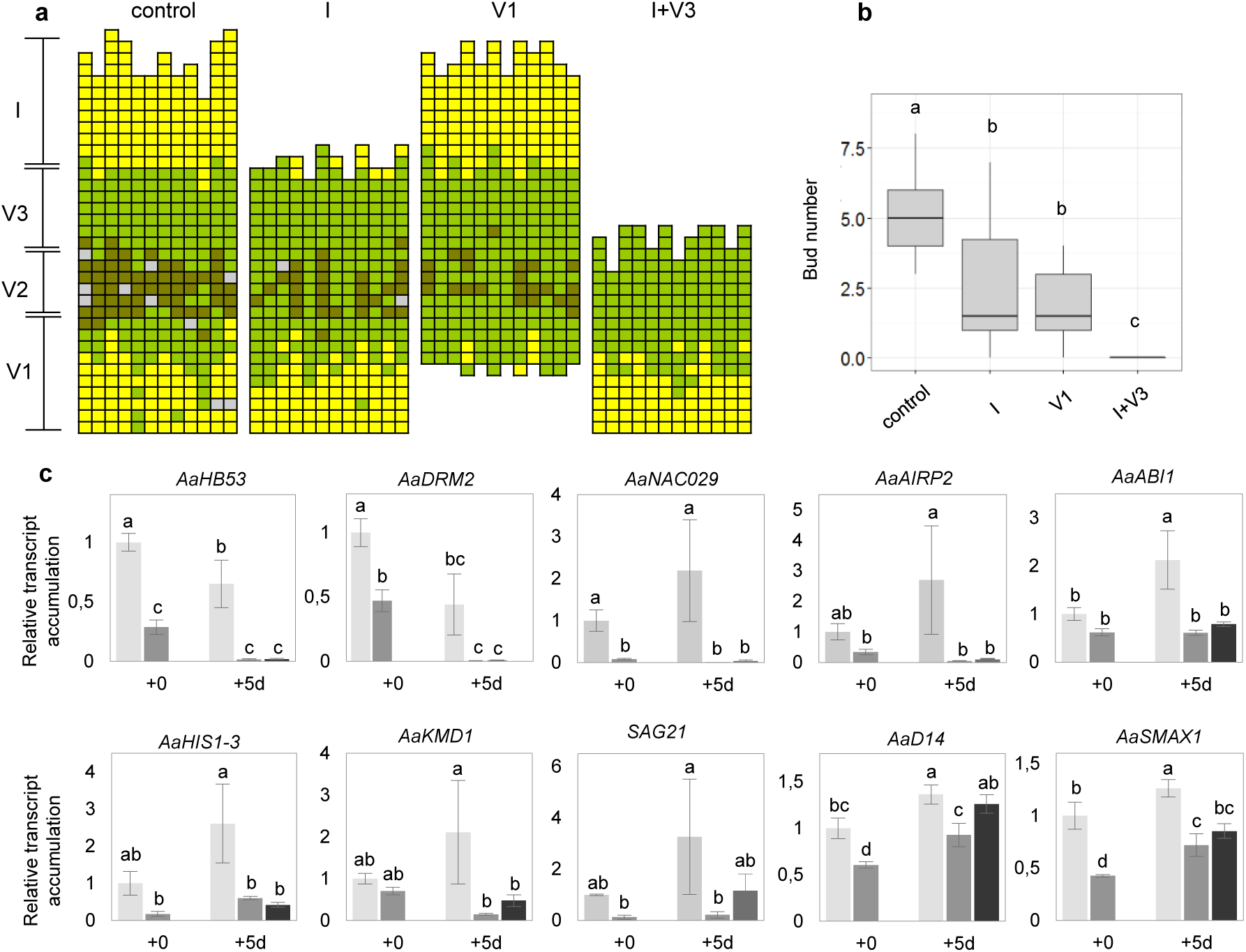
Dormancy of V2 buds is regulated by an apical signal after vernalization. **a**, Analysis of branch formation after excision of axillary branches or buds belonging to different zones in vernalized plants. *Control* indicates intact plants, *I* indicates plants in which the inflorescence buds have been dissected, *V1* indicates plants in which V1 branches have been removed, *I+V3* indicates plants in which the inflorescence and V3 buds have been dissected. As in Fig. 1, each column represents a single plant and each square within a column indicates an individual leaf axil. The bottom row represents the oldest leaf axil. *grey* denotes an empty leaf axil, *brown* denotes a leaf axil with a branch smaller than 0.5 cm, *green* denotes the presence of a vegetative axillary branch and *yellow* denotes the presence of a flowering branch. (*n*= 10 or 12). **b**, Number of buds per plant after the excision of buds or branches in different zones. **c**, Relative transcript accumulation of *AaHB53*, *AaDRM2, AaNAC029, AaAIRP2*, *AaABI1, AaHIS1-3*, *AaKMD1* and *AaSAG21, AaD14* and *AaSMAX1* in V2 (light grey) and V3 buds (dark grey) at end of vernalization (+0) and five days after vernalization (+5d) in control (V2) and decapitated plants (V2 decapitated, black). Expression levels of all genes was normalized with *AaPP2A* and *AaUBI10*. Letters show significant differences between conditions (*P*<0.05, *n*=3) using ANOVA followed by pairwise multiple comparison using Tukey. Errors indicate SD.

### Inhibition of V2 buds is controlled by paradormancy after vernalization and correlates with the increase of auxin in the stem

To identify whether the outgrowth of V2 buds is determined by other parts of the plant or by environmental conditions, we performed a series of decapitation and excision experiments (Fig. 3a). We first addressed the bud behavior in the V2 zone after vernalization by excising buds or branches at the end of vernalization. Excision of inflorescence buds (I) or V1 branches separately reduced the number of buds observed in the V2 zone (nodes 11–16; Fig. 3a, b). The biggest effect on bud outgrowth in the V2 zone was observed when we excised both inflorescence and V3 buds (I+V3) together. Removal of V3 buds alone was only feasible 2 weeks after vernalization when the V3 branches were already expanding and only had a slight effect on the V2 zone (Supplementary Fig. 6). These results suggest that the outgrowth of buds within the V2 zone is influenced by other parts of the plant after vernalization. We also tested the effect of decapitation in plants exposed to different durations of vernalization (Supplementary Fig. 7a). In all treatments, the branches in the V2 zone responded to decapitation and were longer compared to non-decapitated controls (Supplementary Fig. 7b). Interestingly, decapitation of plants vernalized for 12 weeks showed the biggest effect compared to the controls suggesting that stable repression of V2 buds occurs only after 12 weeks of vernalization (Supplementary Fig. 7b). These results suggest that V2 buds are not endodormant during vernalization. To assess the effect of the vernalization treatment on bud outgrowth we decapitated plants and subsequently exposed them to 12 weeks of vernalization (Supplementary Fig. 7c). Decapitated and intact vernalized plants showed a similar number of buds in nodes 0–11 suggesting that cold during vernalization imposes an ecodormant state in the V2 buds (Supplementary Fig. 7d,e). Overall, these results suggest that V2 buds are ecodormant during vernalization and paradormant after vernalization.

To link these results to our transcriptome studies we tested the expression patterns of a set of identified genes after decapitation (Fig. 3c). Most genes tested showed higher transcript accumulation in V2 buds compared to V3 buds at the end and/or after vernalization and their transcript accumulation was reduced in decapitated plants (Fig. 3c). Since auxin is a major regulator of apical dominance, we measured auxin response in the *DR5::GUS* lines after decapitation. At the end of vernalization, stems within the V2 zone did not show strong GUS staining (Fig. 4a,d). A strong GUS signal was detected within 1 week after vernalization in the epidermis and in the vasculature of V2 stems and was reduced in decapitated plants (Fig. 4c,f, Supplementary Fig. 8). We also measured the levels of endogenous IAA in V2 stems of intact plants during the *A. alpina* life cycle. Similar to *DR5::GUS* results, IAA levels in V2 stems were reduced during vernalization and transiently increased again after vernalization (Fig. 4g). Interestingly, IAA levels at 2 weeks after vernalization were very high, which correlated with high IAA levels in the stems of the rapidly growing tissues (inflorescence and V3 branches) above the V2 zone (Fig. 4h,i). Decapitation at the end of vernalization also resulted in a decrease of the endogenous IAA levels in V2 stems (Fig. 4j). This result suggests that the faster outgrowth of the inflorescence and V3 branches after vernalization induces a higher IAA in the V2 stem. To assess the role of auxin and auxin transport in the inhibition of the V2 buds we applied the synthetic auxin NAA and the transport inhibitor NPA in vernalized plants before transferring them to normal greenhouse conditions. NAA treatment increased the number of inhibited buds (Fig. 4k,l) whereas NPA treatment reduced the number of inhibited buds in the V2 zone compared to mock-treated plants (Fig. 4l). In addition, NPA strongly impaired the development of the inflorescence and V1 branches (Fig. 4k). These results confirmed the importance of auxin levels and auxin transport for the inhibition of the buds in the V2 zone after vernalization. We subsequently measured the IAA transport capacity using acropetal ^3^H-IAA treatment in excised V2 stem segments from 8-week-old plants in LDs (8wLD), plants vernalized for 12 weeks (+0) and 5 days after vernalization (+5d). ^3^H-IAA levels in V2 stems were higher before and after vernalization, compared to at the end of vernalization (Fig. 4m,n). Our results suggest that vernalization leads to a decrease in auxin transport in V2 stems (Fig. 4o). Altogether we conclude that the outgrowth of the inflorescence and the vegetative branches after vernalization correlates with an enhancement of endogenous IAA levels, auxin response and transport in the V2 zone after vernalization which may stably repress the outgrowth of V2 buds (Fig. 4o). During vernalization, although auxin transport is low, the development of V2 axillary buds is inhibited (Fig. 4o).

**Fig. 4:**
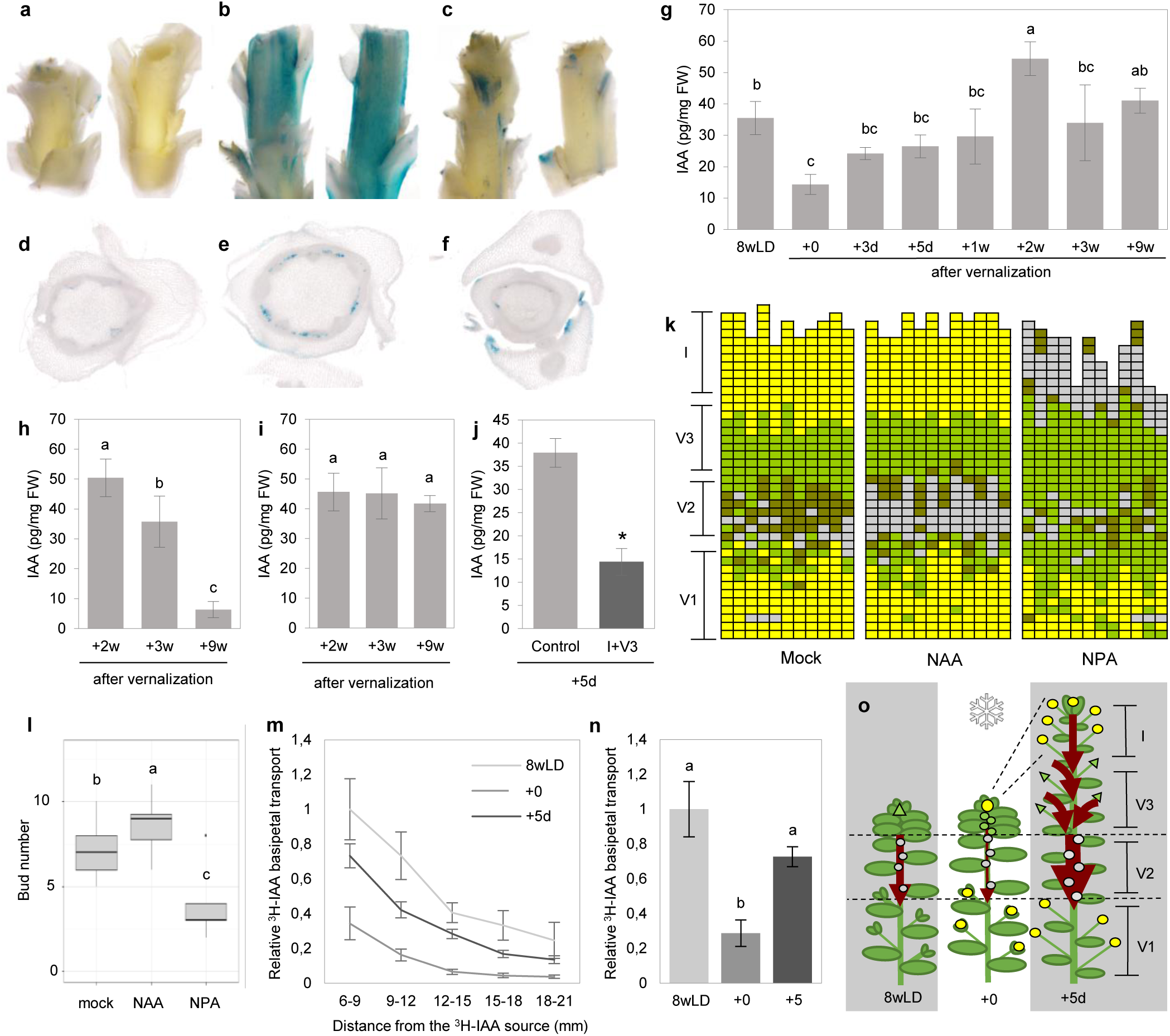
Endogenous IAA levels and auxin response in the stem within the dormant bud zone increase transiently after vernalization. **a-f**, GUS activity in stems within the V2 zone **a-c**, and transversal sections of V2 stem (**d-f**) at end of vernalization (**a,d**) and five days after vernalization (**b,c,e,f**) in control (**b,e**) and decapitated (**c,f**) *DR5::GUS* plants. **g-j**, IAA level in pg/mg of Fresh Weight (FW) before vernalization (8wLD), end of vernalization (+0), 3 and 5 days after vernalization (+3d and +5d), and 1, 2, 3 and 9 weeks after vernalization (+1w, +2w, +3w, and +9w) measured at the V2 stems (**g**) at the base of the inflorescence stem (**h**), at the base of the V3 axillary vegetative branches (**i**) and at V2 stems 5 days after decapitation (**j**). Letters indicate significant differences between conditions (*P*<0.05, *n*=3) using ANOVA followed by pairwise multiple comparison using Tukey. Asterisks in (**j**) indicate significant differences between control and decapitated plants using student’s *t* test (*P*<0.05, *n*=3). Errors indicate SD. **k-l**, Analysis of branch formation in *A. alpina* plants vernalized for 12 weeks and subsequently sprayed with mock or 100μM NAA or 100μM NPA at the end of vernalization and one week after vernalization. Plants were scored 5 weeks after vernalization. As in Fig. 1, each column represents a single plant and each square within a column indicates an individual leaf axil. The bottom row represents the oldest leaf axil. *grey* denotes an empty leaf axil, *brown* denotes a leaf axil with a branch smaller than 0.5 cm, *green* denotes the presence of a vegetative axillary branch and *yellow* denotes the presence of a flowering branch. (**l**) Number of buds per plant (represented with brown and grey boxes) after mock, NAA or NPA treatment. Letters indicate significant differences between conditions (*P*<0.05, *n*=12) using ANOVA followed by pairwise multiple comparison using Tukey. **m-n**, IAA transport capacity in V2 stems in 8 week old plants grown in LD (8wLD), at end of vernalization (+0) and 5 days after vernalization (+5d). ^3^H-IAA measured in (**m**) mm of stem from the ^3^H-IAA source and (**n**) total ^3^H-IAA in stem. Errors indicate SE. **o**, Working model illustrating the reduction of the auxin transport in the V2 zone compared to before (8wLD), at the end of (+0) and 5 days (+5d) after vernalization. Yellow circle denotes flowering of the main or side shoots, grey circle indicates the presence of dormant buds, green triangle the presence of vegetative growth. Red arrow indicates auxin flow in stem.

### AaBRC1 ensures the maintenance of the dormant V2 buds after vernalization

We subsequently tested the expression of genes differentially expressed in our transcriptome analysis on V2 buds across the *A. alpina* life cycle, before, at the end and after vernalization. In addition to our candidate genes we followed the expression patterns of the *A. alpina* homolog of *BRC1* (*AaBRC1*) and its paralog *BRC2* (*AaBRC2*), as in *A. thaliana* the majority of our candidate genes are misexpressed in the *brc1* mutant (Supplementary Fig. 9a)^46^. Compared to before vernalization, *AaBRC1* transcript levels in the V2 buds were increased after prolonged exposure to cold and were maintained in buds harvested 5 days after vernalization (Fig. 5a). Transcript accumulation of the *HB53* homolog (*AaHB53*), a direct target of BRC1^47^, and other BRC1-downstream genes also showed an increase of expression in vernalization (Fig. 5a). Interestingly, transcript accumulation of *AaHIS1-3* only increased after vernalization, suggesting a BRC1-independent regulation of bud dormancy in *A. alpina* (Fig. 5a). We also created transgenic lines with reduced expression of *AaBRC1*. Plants of *35S:AaBRC1dsRNAi* lines 1 and 2 showed significant downregulation of *AaBRC1* expression whereas line 3 did not (Supplementary Fig. 9b). We characterized the *35S:AaBRC1dsRNAi* lines in the same conditions (before, at the end and after vernalization). Before and at the end of vernalization we observed no major difference in branch number and branch length (Fig. 5b,c, Supplementary Fig. 10a,b). Transgenic lines also flowered with a similar total number of leaves compared to control plants (Supplementary Fig. 10 d-h). We observed a phenotype only after vernalization, and nodes 12-16 corresponding to the V2 zone in *35S:AaBRC1dsRNAi* lines 1 and 2 were occupied by a branch (Fig. 5b,e,f, Supplementary Fig. 10c). In these lines, transcript levels of *AaHB53* were reduced in V2 buds of *35S:AaBRC1dsRNAi* lines 1 and 2 at the end and after vernalization but not before vernalization (Supplementary Fig. 10). Interestingly, the transcript accumulation of *AaHIS1-3* and *AaAIRP2* was not misregulated in the *35S:AaBRC1dsRNAi* lines in all developmental stages (Supplementary Fig. 10). These results suggest that the activity of V2 buds is regulated by AaBRC1 after vernalization, although an AaBRC1-independent pathway might exist.

**Fig. 5:**
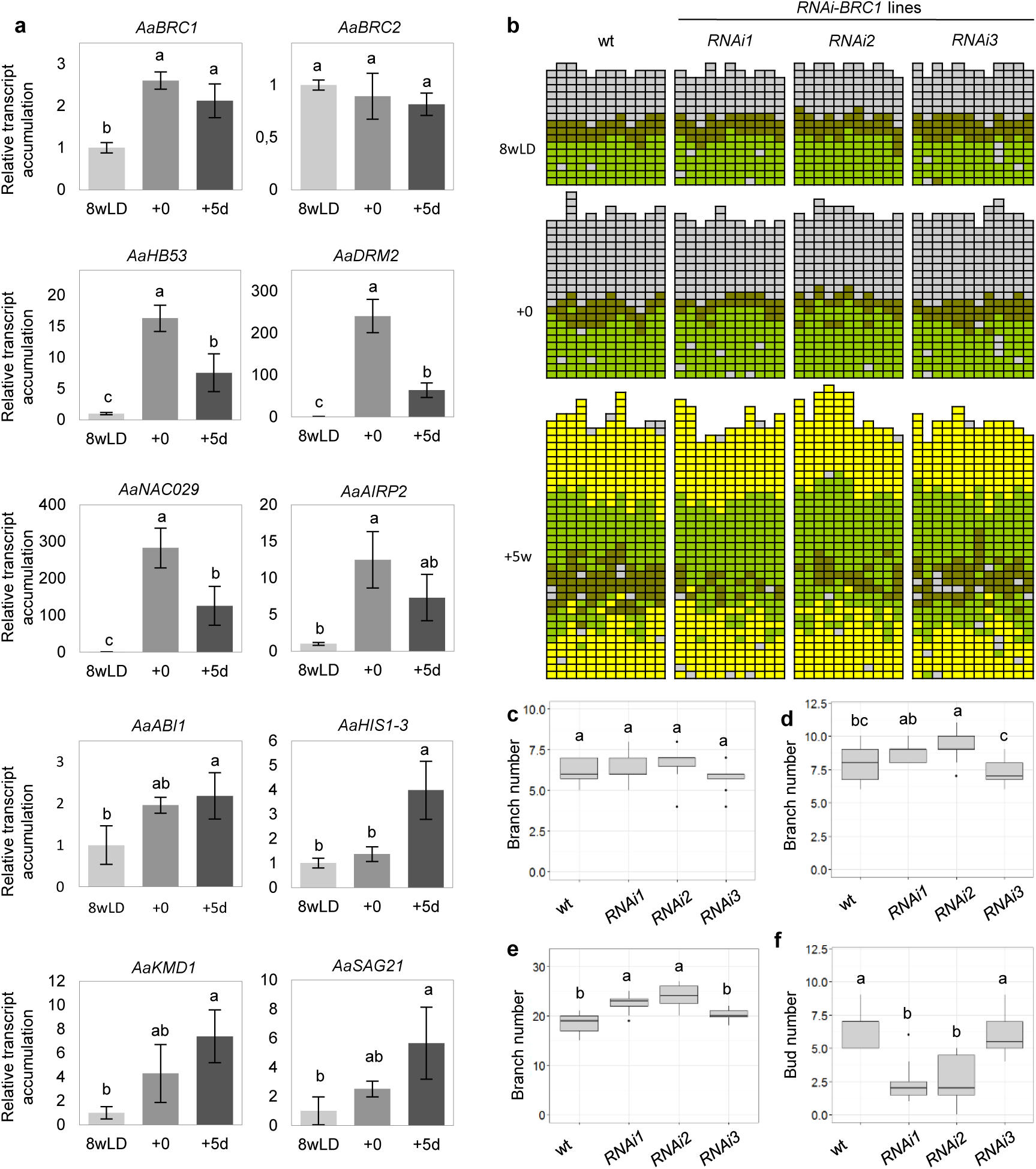
AaBRC1 ensures the maintenance of the dormant V2 buds after vernalization. **a**, Relative transcript accumulation of *AaBRC1, AaBRC2, AaHB53*, *AaDRM2, AaNAC029, AaAIRP2*, *AaABI1, AaHIS1-3*, *AaKMD1* and *AaSAG21* in 8-week-old plants grown in LD (8wLD; light grey), at the end of vernalization (+0; grey) and five days after vernalization (+5d, dark grey). Transcript levels of all genes are normalized with *AaPP2A* and *AaUBI10*. (*n*=3). Errors indicate SD. **b**, Analysis of branch formation in wild-type (wt) plants and in *35S:AaBRC1dsRNAi* lines 1 to 3 in 8-week-old plants grown in LD (8wLD), at the end of vernalization (+0) and five weeks after vernalization (+5w). As in Fig. 1, each column represents a single plant and each square within a column indicates an individual leaf axil. The bottom row represents the oldest leaf axil. *grey* denotes an empty leaf axil, *brown* denotes a leaf axil with a branch smaller than 0.5 cm, *green* denotes the presence of a vegetative axillary branch and *yellow* denotes the presence of a flowering branch. *n*=12. **c-e**, Branch number per plant in wt plants and in *35S:AaBRC1dsRNAi* lines 1 to 3 measured in 8-week-old plants grown in LDs (**c**), at the end of the vernalization period (**d**) and 5 weeks after vernalization (**e**). **f**, Bud number per plant 5 weeks after vernalization. Letters indicate significant differences between conditions (*P*<0.05) using ANOVA followed by pairwise multiple comparison using Tukey.

## DISCUSSION

The induction of flowering in temperate perennials is uncoupled from anthesis so that the flowering process takes place for several years^8,10,48,49^. In the alpine perennial *A. alpina*, flower buds are formed during vernalization and emerge when plants experience favorable environmental conditions. Despite the differences in flowering behavior, similar principles of bud activation described in *A. thaliana* also apply in *A. alpina*. For example, axillary buds close to the shoot apical meristem are temporarily inhibited during vegetative development and the initiation of flowering results in the activation of the upper most axillary buds^13^. However, floral development prior to anthesis is the key for the establishment of a zone of dormant axillary buds. During vernalization, flowering in the main shoot apex always correlates with the presence of V3 buds^19^. Our transcriptome analysis suggests that at the end of vernalization V3 buds are not dormant and might not be dormant throughout the vernalization period. In tulip bulbs, which show a similar pattern of axillary bud activity, the axillary buds located close to the flowering shoot apex never arrest growth^50^. However, exposure to warm temperatures is still required to achieve sustained growth of the V3 buds. This suggests that, in contrast to other species, the phases of bud activation and sustained growth are temporarily separated in *A. alpina*. The link between flowering and paradormancy has been demonstrated in *A. thaliana* and rice with the flowering time regulator FLOWERING LOCUS T/Heading date 3a acting as the systemic signal for flowering and axillary bud activation^51–53^. In *A. alpina*, the *pep1-1* mutant has a clear branching phenotype suggesting that PEP1 might regulate the crosstalk between flowering and bud activity. Other MADS box proteins have been reported in woody perennials to regulate endodormancy and the requirement of prolonged exposure to cold to break dormancy^54^.

Buds in the V2 zone probably transition between different forms of dormancy. Before vernalization, V2 buds are only temporarily dormant due to apical dominance and during exposure to vernalization become ecodormant. This is in contrast to studies in woody perennials, in which buds enter a deeper form of dormancy – endodormancy – during the winter. From the transcript levels of dormancy marker genes we can conclude that the dormancy status of V2 buds is enhanced during vernalization. We detected genes associated with cell cycle and cell division, ABA biosynthesis and signaling to be differentially expressed between dormant (V2) and non-dormant buds (V3). The expression of many of the identified genes in our study has been previously reported to be dependent on BRC1^46^ and is induced under carbon limiting conditions and bud dormancy in *A. thaliana*, grapevine and poplar buds^39^. In our system, the enhancement of bud dormancy during cold correlates with the development of the inflorescence meristem^10,19^. This suggests that sugar demand for inflorescence development during vernalization might be responsible for the carbon starvation response observed in axillary buds. The timing of bud initiation, whether it occurs before or during vernalization might also be an important factor as the carbon starvation response might explain the dormancy status of V2, but not the activation of V3 buds. The link between flowering and bud activity has also been explained by the release from apical dominance due to changes in polar auxin transport^13^. At the end of vernalization, the low IAA levels indicate either that after prolonged exposure to cold the levels of auxin transported basipetally is diminished probably due to a generalized slowdown of growth or that cold influences the auxin metabolism. After the return to warm temperatures, V2 buds once more enter a paradormant state being dominated by the inflorescence and V3 or V1 branches. The transition from the ecodormant back to the paradormant state involves the enhancement of basipetal auxin transport after vernalization and increased activity of AaBRC1.

Overall, we conclude that *A. alpina* is a good model system to follow different stages of dormancy across the perennial life cycle and to dissect the phases of bud outgrowth, release of dormancy and sustained branch growth.

## Methods

### Plant material, growth conditions and phenotyping

All our experiments were performed with the *Arabis alpina* accession Pajares or the *pep1-1* mutant described by Wang et al.^19^. Seeds were stratified on humidified paper at 4°C in the dark for 4 days prior to germination on soil under temperatures ranging from 20 °C during the day to 18 °C during the night in a long day (LD, 16 hours light, 8 hours dark) greenhouse. After 8 weeks of growth, vernalization treatments were carried out in a short day (SD, 8 hours light, 16 hours dark) growth chamber at 4°C for different durations of vernalization (depending on the experiment) before the plants were moved back to a LD (16 hours light, 8 hours dark) greenhouse at 20°C. For the juvenility experiment plants were grown for 3 weeks (juvenile) or 5 weeks (adult) before being vernalized. For 1-naphthaleneacetic acid (NAA) and 1-N-naphthylphthalamic acid (NPA) treatments at the end of vernalization and one week after vernalization, plants were sprayed with 100 µM NAA (Sigma-Aldrich), 100 µM NPA (Chem Service) or DMSO as a mock treatment with 0.2% (v/v) Tween-20 immediately after being transferred back to a LD greenhouse. Excision of inflorescence and/or V3 buds at the end of vernalization was performed under a stereomicroscope by removing the eight nodes below of the lowest flowering bud. In the simple decapitation method (Supplementary Fig. 7), the apex was cut off directly above the point where no stem elongation was observed, corresponding to nodes 11–14 within the V2 zone. The shoot architecture at different time points was recorded by observing the fate and the length of the branch at each node of the plant, and the number and type of branches per zone. All experiments were performed with 10 to 12 plants.

### Plasmid constructs

The *DR5::GUS* fragment (excised from plasmid *DR5::GUS,* kindly provided by Tom Guilfoyle) was introduced into the GATEWAY-compatible pEarleyGate 301 plasmid containing the BASTA resistance gene using site-directed recombination. For the *35S:AaBRC1dsRNAi* constructs, three cDNA fragments of *AaBRC1* (Fragment 1-3; see Supplementary Table 5) were amplified and introduced into the GATEWAY-compatible pJawohl-8-RNAi plasmid using site-directed recombination. The names of the *A. alpina 35S:AaBRC1dsRNAi* lines correspond to the fragments introduced. For each construct, homozygous transgenic *A. alpina* lines carrying single-copy transgenes were generated using the floral dip method^55^.

### *GUS* staining

For GUS staining assays, stems within the V2 zone below the point where no stem elongation was observed were harvested and leaf axils carrying V2 and V3 buds of *DR5::GUS* plants were excised and placed directly into 90% ice-cold acetone and incubated for 1 h on ice. V3 buds were identified as the 8 nodes directly below the flowering buds. V2 buds were identified below the V3 buds where no stem elongation was observed. The samples were washed in 50 mM phosphate buffer (pH 7.0) and submerged in 2.5 mL GUS staining solution under moderate vacuum for 20 min^56^. After incubation at 37°C in the dark for a maximum of 16 h, chlorophyll was removed by transferring the samples through an ethanol series. GUS activity was observed in whole stem tissues, transverse stem sections or longitudinal leaf axil sections. We prepared 50–60 µm sections on a Leica VT1000S vibratome from samples immobilized on 6% (w/v) agarose. Representative photographs from two different biological experiments were taken using the stereomicroscope Nikon SMZ18 and Nikon Digital Sight camera (DS-Fi2) for whole stem segments, and the Zeiss Axio Imager microscope with the Zeiss Axio Cam 105 color camera for cuttings.

### RNA extraction and cDNA synthesis

For RNA-Seq transcript profiling and quantitative RT-PCR analysis, V2 and V3 buds were specifically harvested under a stereomicroscope immediately after vernalization and 5 days after vernalization. In all experiments, V2 buds corresponded to the leaf axils 16–19, and V3 buds corresponded to the leaf axils 23–26. For quantification of the *GUS* expression of the *DR5::GUS* lines, the stem was cut off directly below the point where stem elongation was observed, axillary buds were removed so that RNA was isolated only from three internodes within the V2 zone. Each experiment comprised three independent biological replicates. Samples were stored at –80°C prior to RNA extraction. Total RNA was extracted using the RNeasy Plant Mini Kit (Qiagen) following the manufacturer’s instructions, followed by a DNase treatment with Ambion DNA free-kit DNase treatment and removal (Invitrogen). Total RNA (2 µg) was used for the synthesis of cDNA by reverse transcription with SuperScript II Reverse Transcriptase (Invitrogen) and oligo dT (18) primers.

### RNA sequencing transcript profiling

Poly(A) RNA enrichment, library preparation and sequencing were carried out at the Max Planck Genome Center, Cologne, Germany (https://mpgc.mpipz.mpg.de/home/) using 1 µg total RNA. Poly(A) RNA was isolated using the NEBNext Poly(A) mRNA Magnetic Isolation Module (New England Biolabs) and the library was prepared using the NEBNext Ultra Directional II RNA Library Prep Kit for Illumina (New England Biolabs). RNA quality and quantity were checked by capillary electrophoresis (TapeStation, Agilent Technologies) and fluorometry (Qubit, Thermo Fisher Scientific) after each step. Sequencing was carried out using a HiSeq3000 (Illumina) with 1 x 150 bp single reads.

Reads derived from the Illumina library were mapped and aligned to the reference genome using HISAT2 followed by assembly and quantification of expression levels in different samples using STRINGTIE. The gene counts of all samples were obtained by using a Python script (http://ccb.jhu.edu/software/stringtie/dl/prepDE.py). The quality of the samples was assessed by producing dispersion plots among replicates. The differentially expressed genes with more than 2-fold change and a corrected p-value below 0.05 were obtained using DESeq2 and selected for further analysis. We focused on genes with a greater than 2-fold change in transcript abundance. The complete transcriptome data set is available as series GSE126944 at the NCBI Gene Expression Omnibus (http://www.ncbi.nlm.nih.gov/geo/). The Database for Annotation, Visualization and Integrated Discovery (DAVID; Huang et al., 2009) was used for Gene Ontology enrichment analysis for biological processes 5, with Benjamini corrected *P*<0.05. Only Arabidopsis annotated orthologues were included for GO analysis. Data were clustered using Cluster 3.0 and visualized using Java TreeView (http://doi.org/10.5281/zenodo.1303402). From the 25,817 genes which showed transcript accumulation in at least one of the conditions tested, 4983 participated in the hierarchical clustering. Only 11% of genes in the cluster did not behave similarly between replicates (Supplementary Table S2; clusters I0 and II0). Venn diagrams were constructed using Venny v2.1 (http://bioinfogp.cnb.csic.es/tools/venny/).

### Quantitative real-time PCR

Three technical replicates were prepared using 26 ng of cDNA for each reaction. The relative gene expression values were based on ΔCt calculations using the mean of the two reference gene expression values, according to Pfaffl et al.,^58^. *AaUBI10* and *AaPP2A* were used for expression data normalization. The ΔCt values were scaled to the average for the control condition. Primers used can be found in Supplementary Table 6.

### Quantification of IAA

For free IAA quantification, V2 stems were cut off directly below the point where stem elongation was observed, axillary buds were removed so that IAA was measured only from three internodes within the V2 zone. Inflorescence (I) and V3 stems 2cm from the base of stems were harvested. Plant material (around 15 mg fresh weight) was purified as previously described in Andersen et al.^59^, and 500 pg ^13^C_6_-IAA internal standard was added to each sample before homogenisation and extraction. Free IAA was quantified in the purified samples using combined gas chromatography - tandem mass spectrometry.

### ^3^H-IAA transport assay

Stem segments (21 mm) from 8-week-old plants (8wLD), from plants at the end of vernalization (+0) and from plants 5 days after vernalization (+5) were cut off directly above the point where no stem elongation was observed. The segments were placed on wet paper and transferred to 30 μl 0.05% MES (pH 5.5–5.7) containing 100 nM ^3^H-IAA (Hartmann Analytic). After incubation for 10 min, the stem segments were transferred to fresh 0.05% MES containing 1 µM IAA for 90 min^60^. Incubation was performed at 4°C for the stem segment of samples harvested at the end of the vernalization. The stems were then cut into 3 mm segments and immersed in Rotiszint eco plus (Roth) for 16 h before the radiolabel was quantified by scintillation for 2 min using a LS6500 Multi-Purpose Scintillation Counter (Beckman Coulter). CPM values were scaled to the average for the 8 week long day plant sample at 6-9 mm or 8 week long day total samples.

### Statistical analysis

Data were processed by analysis of variance (ANOVA) followed by a post hoc test for pairwise multiple comparisons using Tukey post hoc test using the *R* platform (http://r-project.org/). Pairwise comparisons were analyzed using Student’s t test.

## Acknowledgements

M.C.A. and K.T. acknowledge support from the Deutsche Forschungsgemeinschaft (DFG, German Research Foundation) under Germany’s Excellence Strategy – EXC 2048/1 – Project ID: 390686111. K.L. acknowledges support from the Knut and Alice Wallenberg Foundation (KAW), the Swedish Research Council (VR) and the Swedish Governmental Agency for Innovation Systems (VINNOVA). The authors would like to thank Roger Granbom, Lisa Clark, Julia Benecke, Vicky Tilmes, Kirstin Krause and Panpan Jiang for technical assistance or for sharing primers and Margaret Kox for the critical reading of the manuscript.

## Contributions

A.V., P.M., K.T. and M.C.A. conceived the study and designed the experiments; A.V., A.R., K.L., U.N., U.P. performed the experiments; A.V., P.M., A.R., and M.C.A. analysed the data; A.V. and M.C.A. wrote the manuscript with the contributions from all authors.

## Supplementary Figures

**Supplementary Fig. 1:**
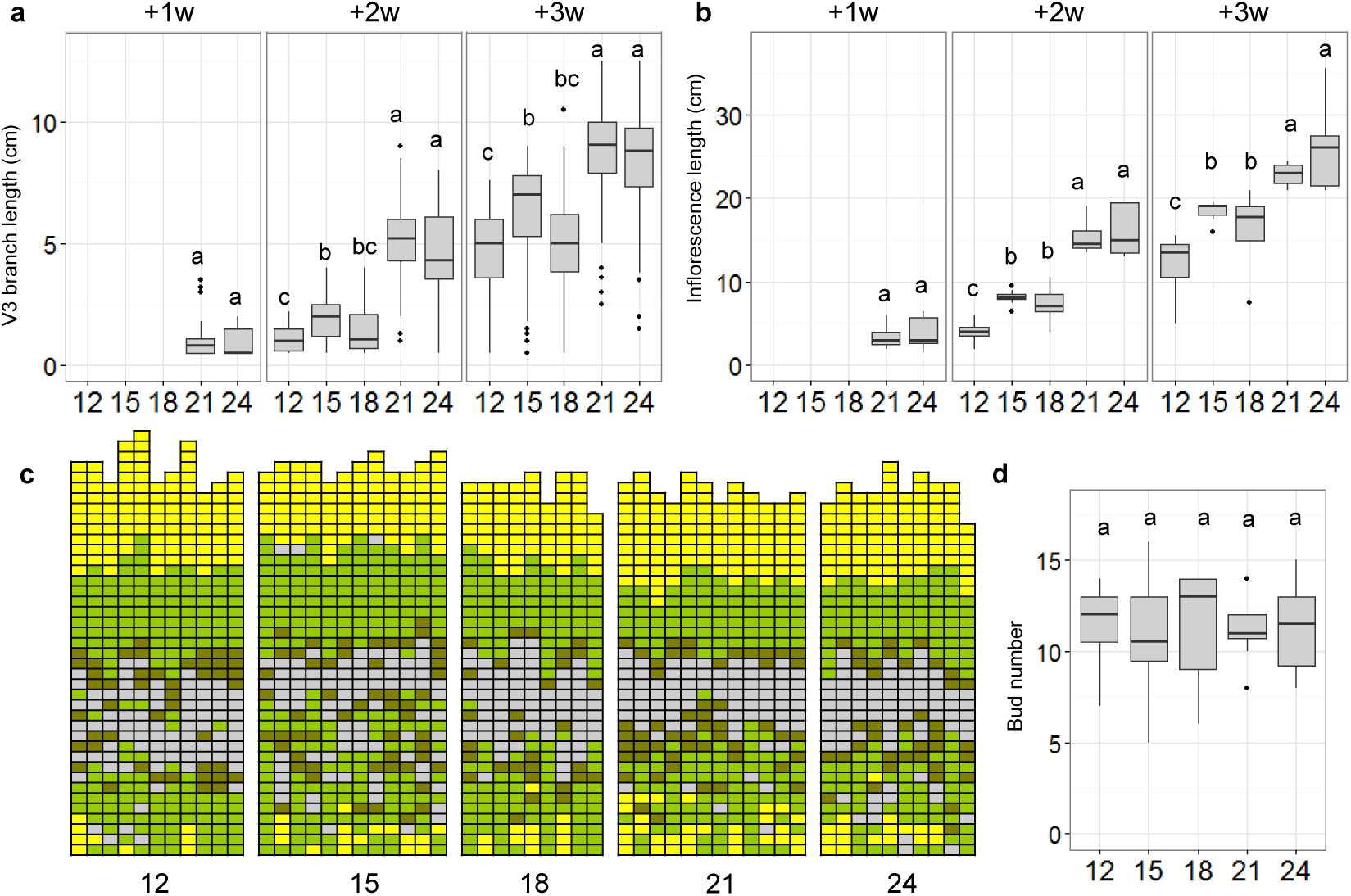
Extended vernalization accelerates the outgrowth of the vegetative branches (V3) and the inflorescence but does not influence the final shoot architecture. **a-b**, Length of the vegetative branches (V3) (**a**) and inflorescence (**b**) at 1, 2 and 3 weeks (+1w, +2w, +3w) after vernalization measured in plants grown for 8 weeks in LDs and vernalized for 12, 15, 18, 21 and 24 weeks. (**c**) Analysis of branch formation in plants vernalized for 12, 15, 18, 21 and 24 weeks, measured 3 weeks after vernalization. As in Fig. 1, each column represents a single plant and each square within a column indicates an individual leaf axil. The bottom row represents the oldest leaf axil. g*rey* denotes an empty leaf axil, *brown* denotes a leaf axil with a branch smaller than 0.5 cm, *green* denotes the presence of a vegetative axillary branch and *yellow* denotes the presence of a flowering branch. **d**, Number of buds per plant (represented with *brown* and *grey* boxes) in plants vernalized for 12, 15, 18, 21 and 24 weeks. Letters show significant differences between conditions (*P*<0.05) using ANOVA followed by pairwise multiple comparison using Tukey. *n*=9-12.

**Supplementary Fig. 2:**
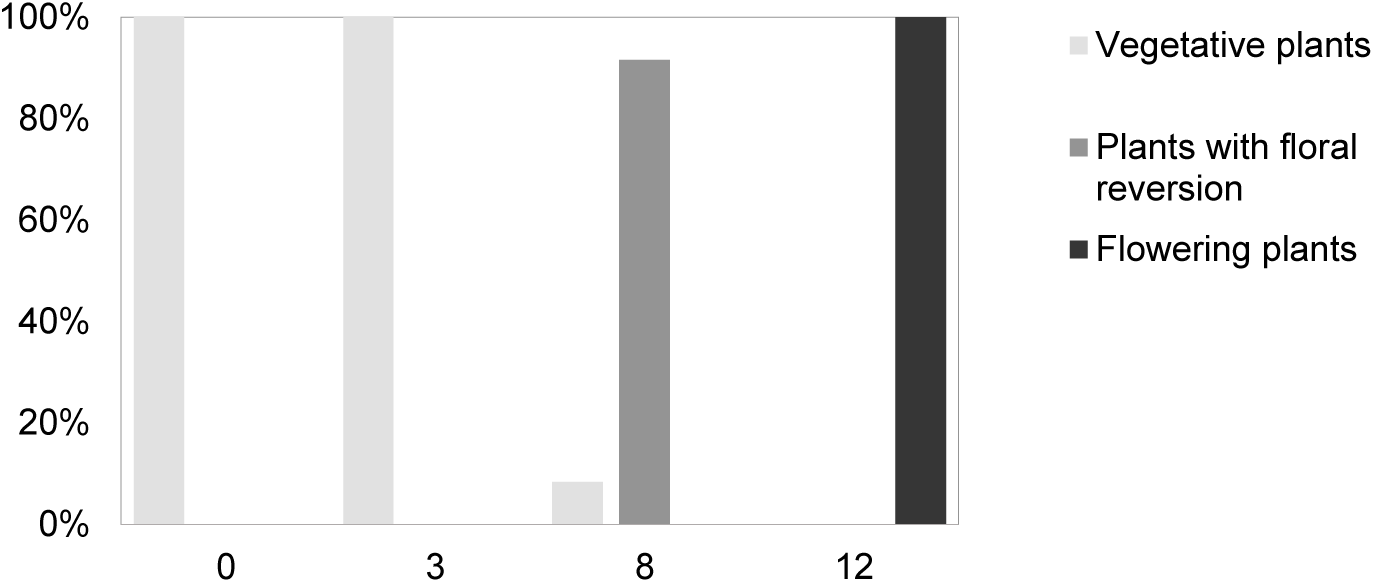
Duration of vernalization influences floral commitment in the shoot apical meristem. Percentage of vegetative plants, plants with floral reversion and flowering plants after different durations of vernalization. Plants were grown for 8 weeks in LDs (0), or vernalized for 3 (3), 8 (8) or 12 (12) weeks. Plants were scored 3 weeks after they were returned to a LD greenhouse, *n*=12.

**Supplementary Fig. 3:**
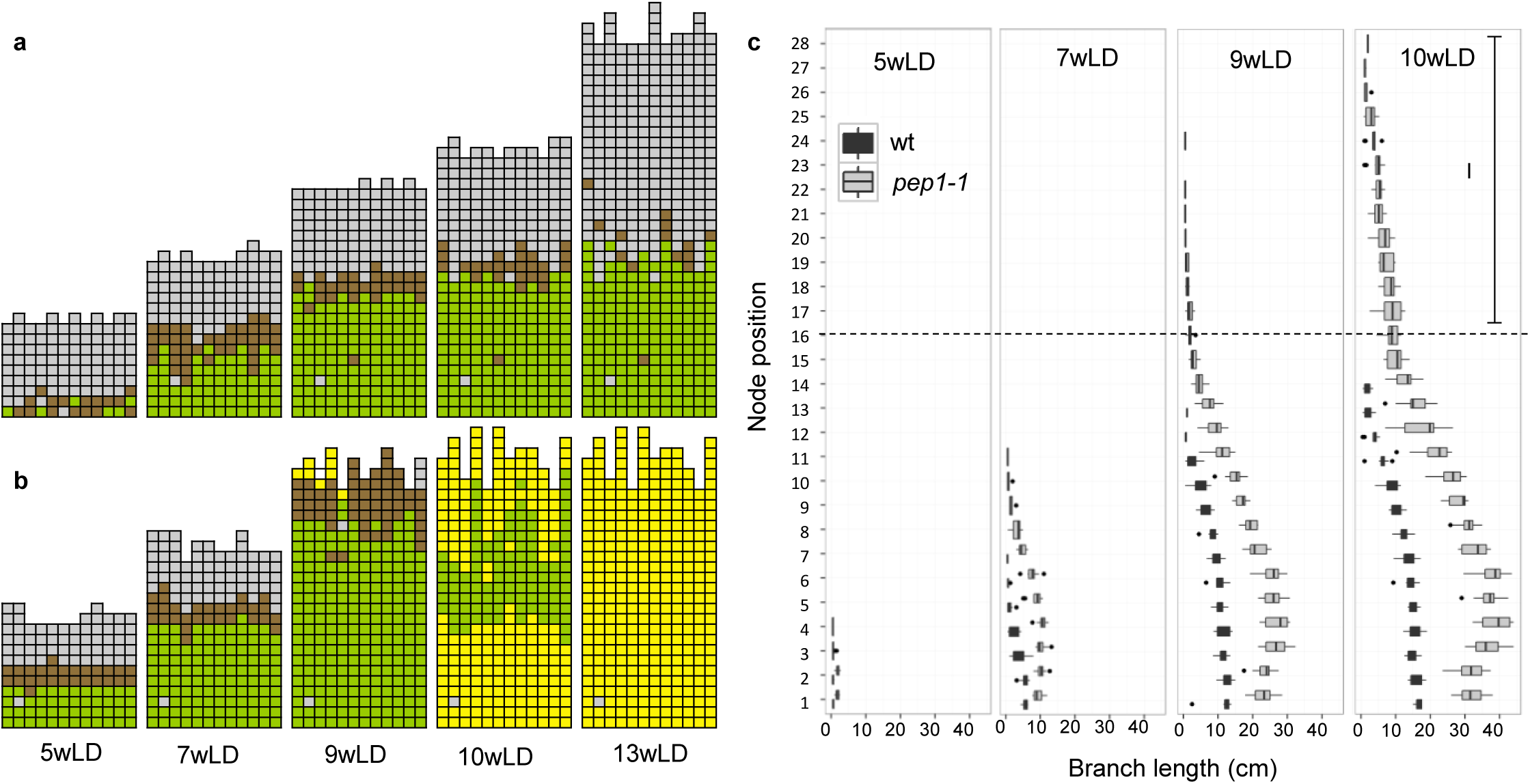
All axils in the *A. alpina pep1-1* mutant develop a flowering axillary branch. **a-b**, Analysis of branch formation in wild type (wt) (**a**) and *pep1-1* mutant (**b**) and **c**, Branch length according to the node position in wt and *pep1-1* mutant growing for 5, 7, 9, 10 and 13 weeks in a long day greenhouse (LD). As in Fig. 1, each column represents a single plant and each square within a column indicates an individual leaf axil. The bottom row represents the oldest leaf axil. g*rey* denotes an empty leaf axil, *brown* denotes a leaf axil with a branch smaller than 0.5 cm, *green* denotes the presence of a vegetative axillary branch and *yellow* denotes the presence of a flowering branch. *I* indicates the inflorescence branches in *pep1-1*. *n*=12.

**Supplementary Fig. 4:**
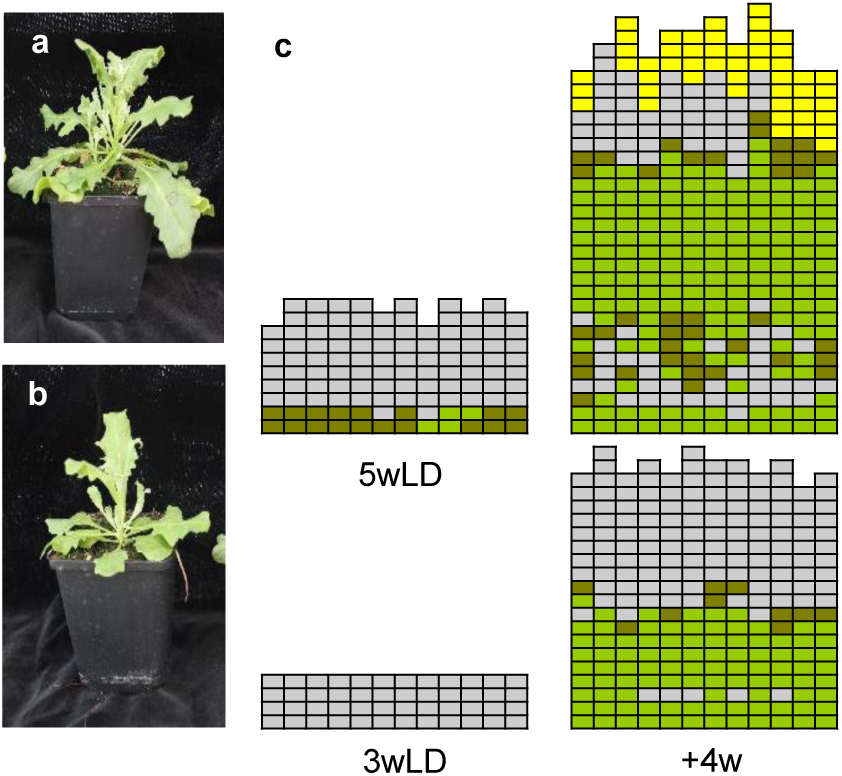
Only flowering plants have a zone with inhibited buds after vernalization. **a-c,** Analysis of branch formation in juvenile and adult plants after vernalization. Plants were grown for 3 weeks (juvenile) or 5 weeks (adult) in long days (LDs) before being vernalized for 12 weeks. Only 5-week-old vernalized plants will initiate flowering during vernalization whereas 3-week-old vernalized plants continue vegetative growth. Pictures of an adult (**a**) and a juvenile (**b**) vernalized plant after being returned for 2 weeks in LDs. For (**c**), similar to Fig. 1, each column represents a single plant and each square within a column indicates an individual leaf axil. The bottom row represents the oldest leaf axil. g*rey* denotes an empty leaf axil, *brown* denotes a leaf axil with a branch smaller than 0.5 cm, *green* denotes the presence of a vegetative axillary branch and *yellow* denotes the presence of a flowering branch.

**Supplementary Fig. 5:**
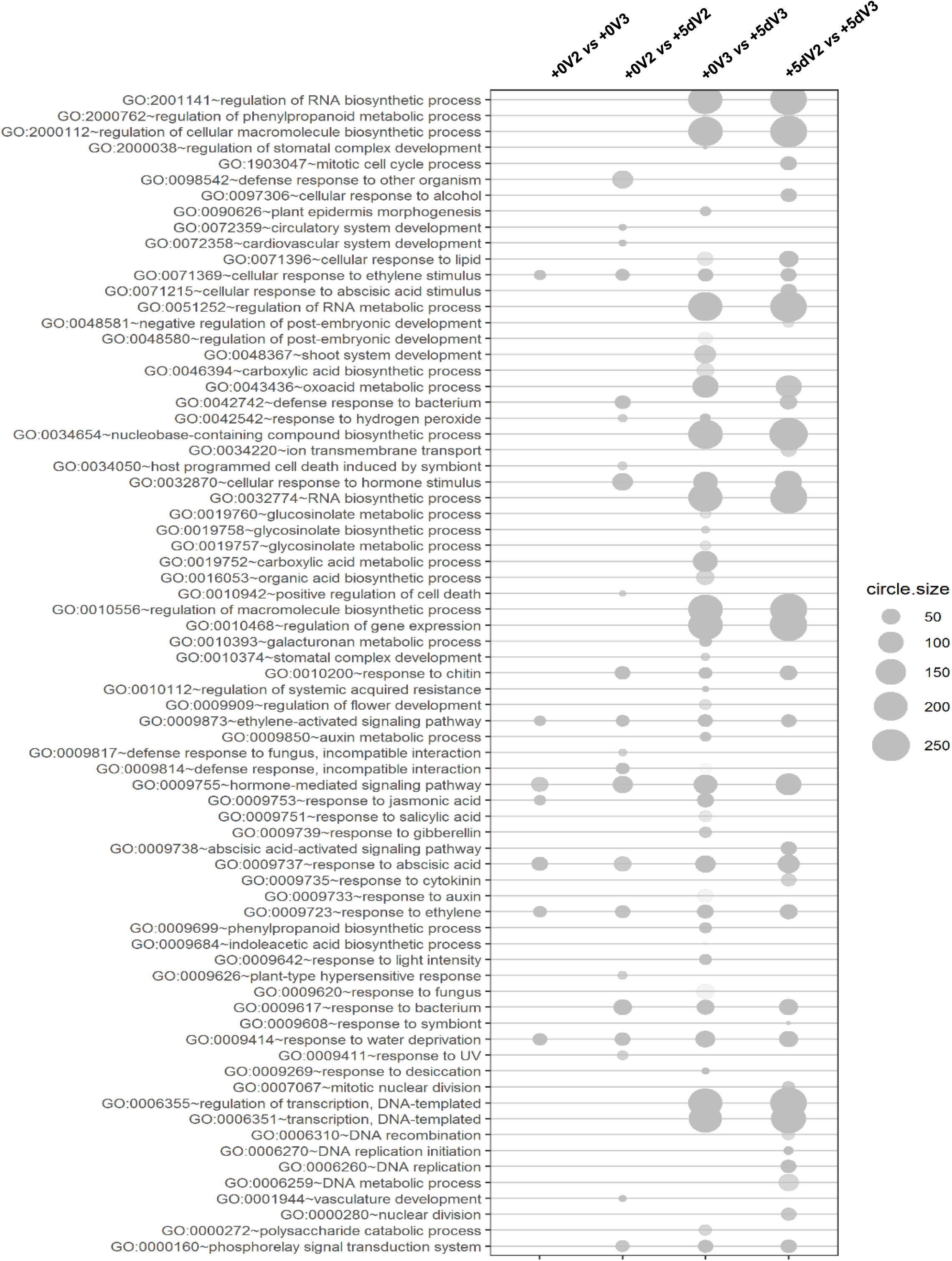
GO enrichment analysis in differentially regulated genes in V2 and V3 buds at the end of vernalization (+0) and five days after vernalization (+5d). Circle size indicats the number of genes in the GO category.

**Supplementary Fig. 6:**
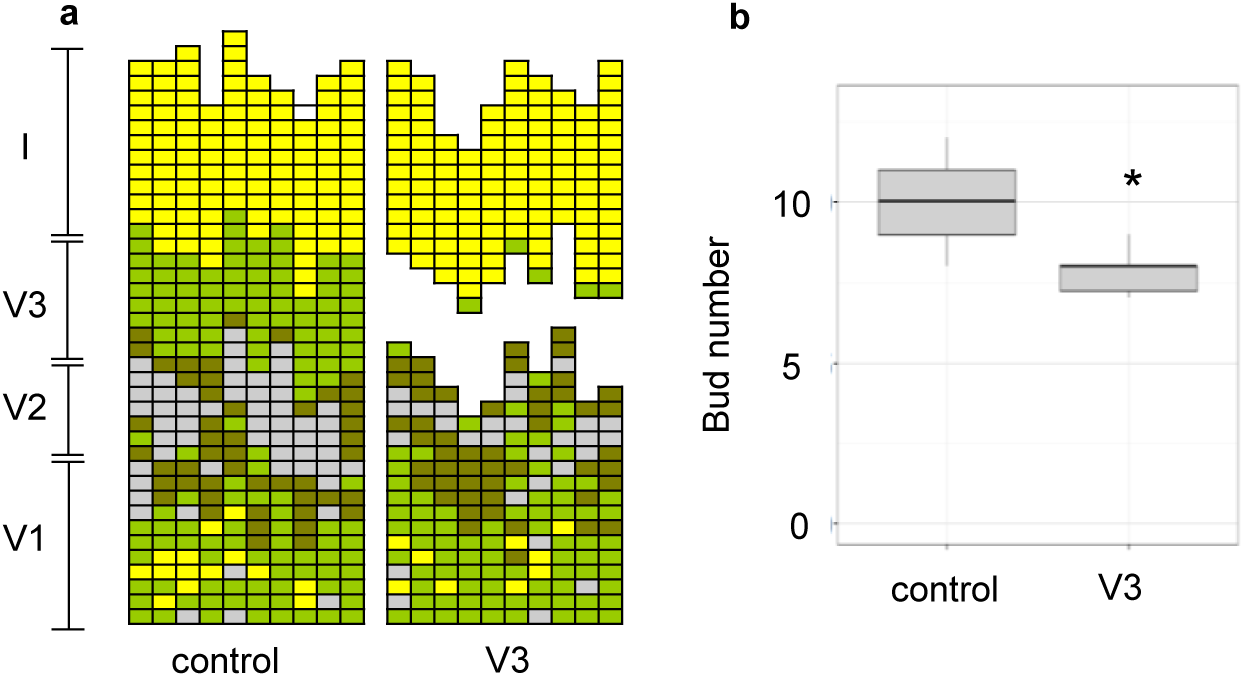
Dormancy of V2 buds is slightly affected by the removal of V3 branches 2 weeks after vernalization. **a**, Analysis of branch formation and **b**, number of buds in plants after removal of axillary vegetative branches in the V3 zone 2 weeks after vernalization. Plants were grown for 8 weeks in long days (LDs) and vernalized for 12 weeks. Scoring of the branching pattern in each node was performed 2 weeks after treatment. As in Fig. 1, each column represents a single plant and each square within a column indicates an individual leaf axil. The bottom row represents the oldest leaf axil. g*rey* denotes an empty leaf axil, *brown* denotes a leaf axil with a branch smaller than 0.5 cm, *green* denotes the presence of a vegetative axillary branch and *yellow* denotes the presence of a flowering branch. Asterisks indicate significant differences using student’s *t* test, *P*<0.05 (*n*= 10).

**Supplementary Fig. 7:**
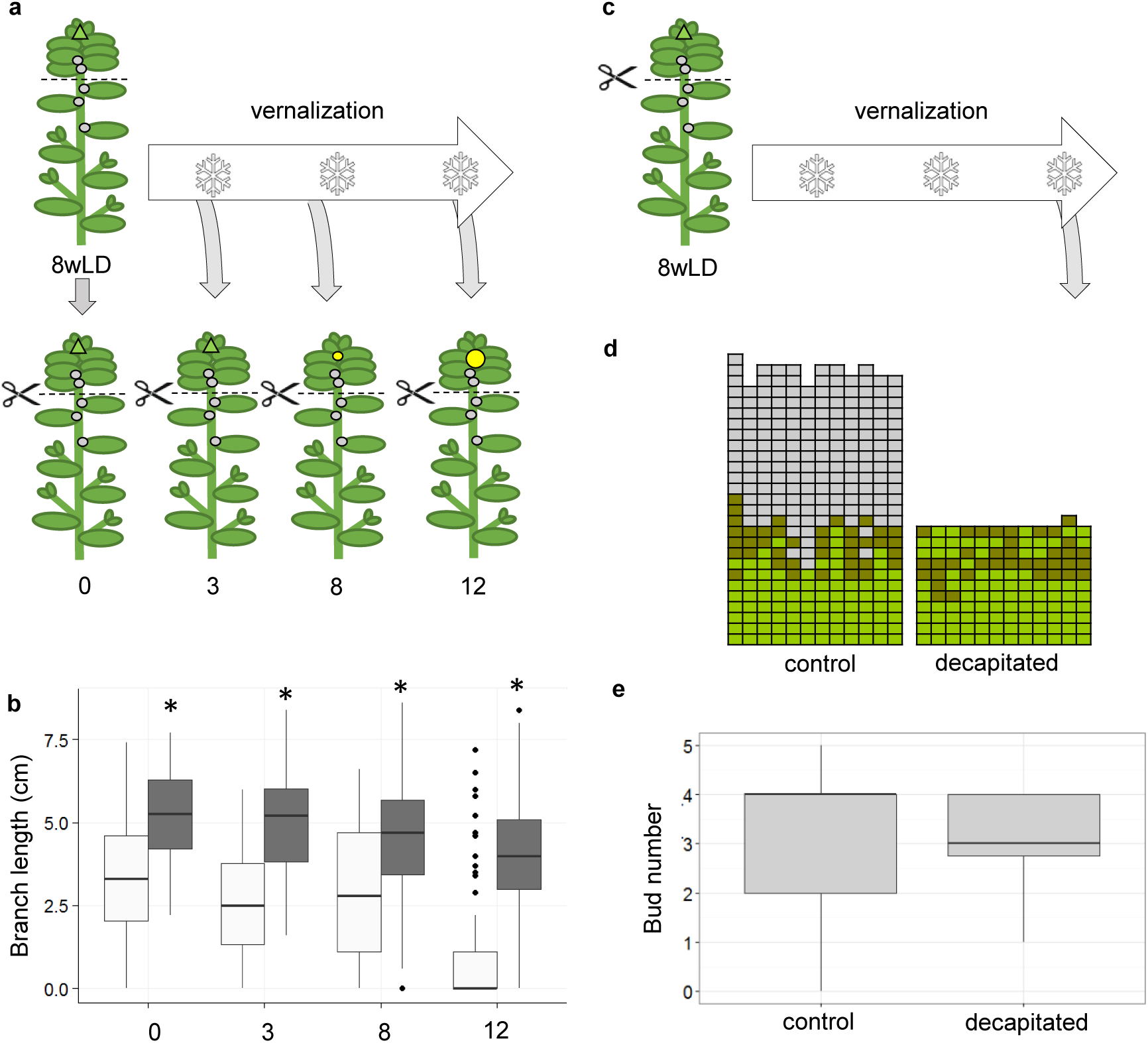
Buds in the V2 zone respond to decapitation before or after different vernalization durations after being returned to LD greenhouse conditions but not when remained in vernalization. **a**, Schematic representation of the experimental design of **b**. Plants were grown for 8 weeks in long days (8wLD) and subsequently vernalized for 0, 3, 8 and 12 weeks. Prior to being returned to warm temperatures, plants were decapitated. **b**, Length of new branches of control (white) or decapitated plants (grey) at 0, 3, 8 and 12 weeks of vernalization. Branch length was scored 2 weeks after decapitation. Control plants are the same as in Fig. 1g and h. **c**, Schematic representation of the experimental design of **d** and **e**. Plants were grown for 8 weeks in long days (8wLD), decapitated and subsequently vernalized for 12 weeks. **d**, Analysis of branch formation in plants at the end of the 12 week vernalization period in control plants and decapitated plants. As in Fig. 1, each column represents a single plant and each square within a column indicates an individual leaf axil. The bottom row represents the oldest leaf axil.g*rey* denotes an empty leaf axil, *brown* denotes a leaf axil with a branch smaller than 0.5 cm and *green* denotes the presence of a vegetative axillary branch. **e**, Number of buds in nodes 1-11 after 12 weeks of vernalization in control plants or decapitated plants. Asterisks indicate significant differences using student’s *t* test *P*<0.05 between control and decapitation. *n*=*12*.

**Supplementary Fig. 8:**
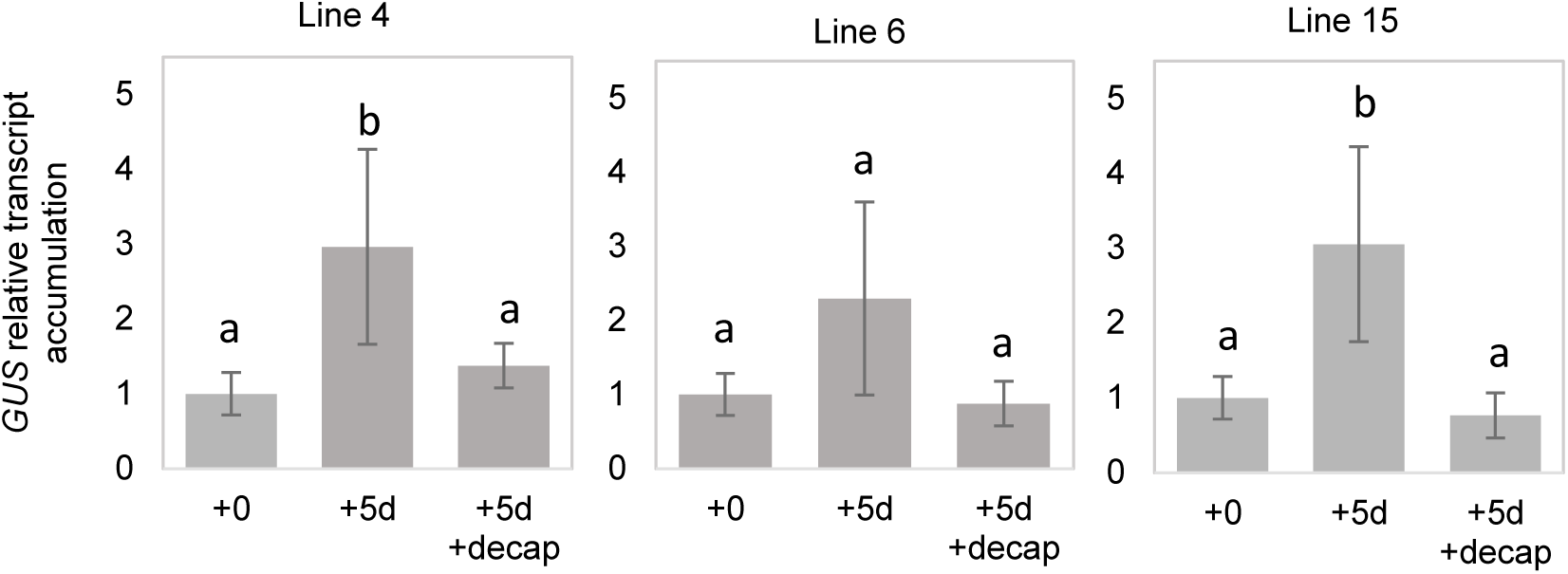
*GUS* transcript accumulation in *DR5::GUS A. alpina* lines 4, 6 and 15. *GUS* transcript accumulation was tested in V2 stems of plants grown for 8 weeks in LDs and vernalized for 12 weeks at the end of vernalization (+0) and 5 days after in control plants (+5d) and decapitated plants (+5d+decap). Samples were normalized with *AaPP2A* and *AaUBI10*. Letters show significant differences between conditions (*P*<0.05, n=3) using ANOVA followed by pairwise multiple comparison using Tukey.

**Supplementary Fig. 9:**
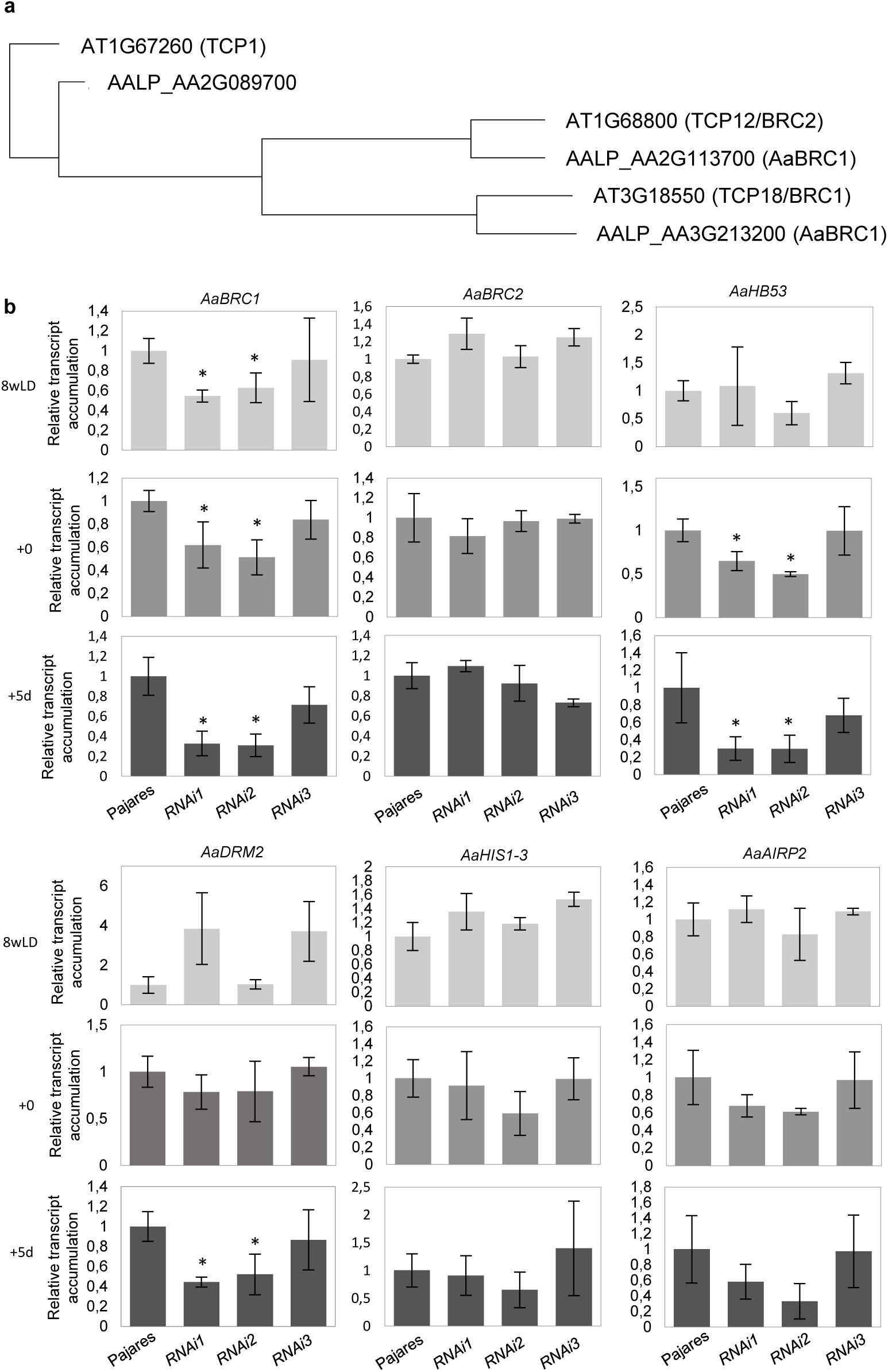
Transcript accumulation of dormancy-associated genes was reduced after vernalization in *35S:AaBRC1dsRNAi* lines. **a,** Phylogenetic tree showing relationship between *A. alpina BRC1* and *BRC2* homologues. **b**, Relative transcript accumulation of *AaBRC1, AaBRC2, AaHB53, AaDRM2, AaHIS1-3* and *AaAIRP2* in V2 buds in 8-week-old plants grown in LD (8wLD; light grey), at end of vernalization (+0; grey) and five days after vernalization (+5d, dark grey) in wt plants and in *35S:AaBRC1dsRNAi* lines 1 to 3. Expression of all genes was normalized with *AaPP2A* and *AaUBI10*. Asterisks indicate significant differences between the tested conditions and the wt using student’s *t* test (*P*<0.05, *n*=3). Errors indicate SD.

**Supplementary Fig. 10:**
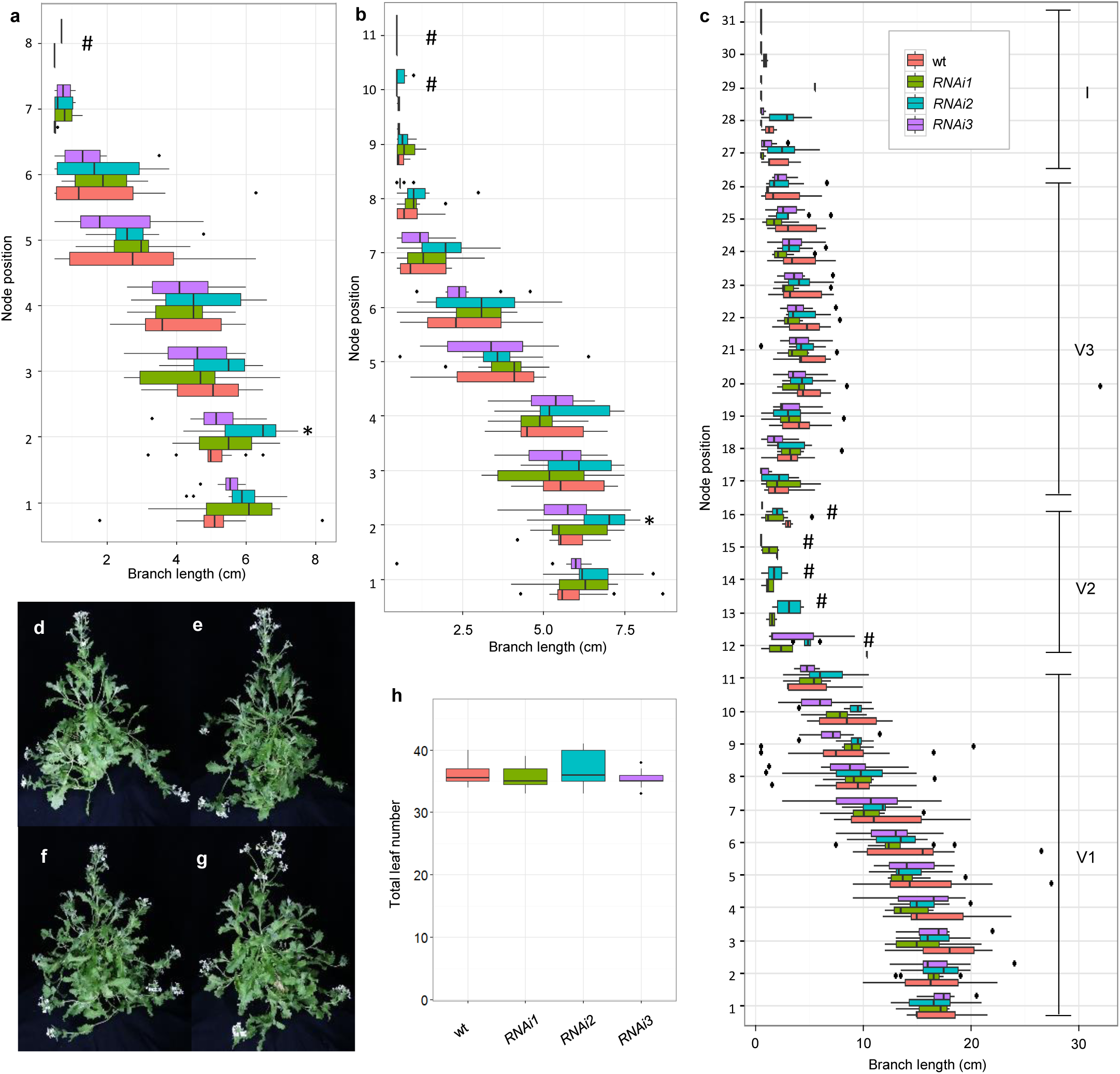
*35S:AaBRC1dsRNAi* lines do not show major differences in branch length and total leaf number. **a-c**, Branch length according to the node position in wt and *35S:AaBRC1dsRNAi* lines 1-3 before- (**a**), at the end- (**b**) and 2 weeks after- (**c**) vernalization. **d-g**, Pictures of the wt (**d**), the *35S:AaBRC1dsRNAi* lines 1 (**e**), line 2 (**f**) and line 3 (**g**) 3 weeks after vernalization. **h**, Total leaf number at flowering in wt and the *35S:AaBRC1dsRNAi* lines 1-3. Asterisks indicate significant differences between the tested condition and the wt using student’s *t* test (*P*<0.05, *n*=10-12). Hashtags indicate nodes where less than 3 branches could be measured for the wt plants.

## Supplementary Tables

**Supplementary Table 1.** List of genes whose transcript levels have been identified to be differentially expressed between V2 and V3 buds at the end of vernalization and 5 days later.

**Supplementary Table 2.** GO enrichment categories identified in genes whose transcript levels have been identified to be differentially expressed between V2 and V3 buds at the end of vernalization and 5 days later.

**Supplementary Table 3.** List of coexpressed clusters obtained after hierarchical clustering of the transcript accumulated in V2 and V3 buds at the end of vernalization and 5 days later.

**Supplementary Table 4.** GO enrichment categories identified in the different coexpressed clusters.

**Supplementary Table 5.** Genes differentially regulated in at least one of the conditions, homologues to *A. thaliana* gene identified as “*bud dormancy*” marker genes in Tarancón et al^39^

**Supplementary Table 6:** Primers used in this article.

